# Red deer *Cervus elaphus* blink more in larger groups

**DOI:** 10.1101/2020.08.11.245837

**Authors:** Zeke W. Rowe, Joseph H. Robins, Sean A. Rands

## Abstract

Most animals need to spend time being vigilant for predators, at the expense of other activities such as foraging. Group-living animals can benefit from the shared vigilance effort of other group members, with individuals reducing personal vigilance effort as group size increases. Behaviours like active scanning or head lifting are usually used to quantify vigilance, but may not be accurate measures as the individual could be conducting them for other purposes. We suggest that measuring an animal’s blinking rate gives a meaningful measure of vigilance: increased blinking implies reduced vigilance, as the animal cannot detect predators when its eyes are closed. We demonstrate that as group size increases in red deer, individuals increase their blink rate, confirming the prediction that vigilance should decrease. Blinking is a simple non-invasive measure, and offers a useful metric for assessing the welfare of animals experiencing an increase in perceived predation risk or other stressors.

## INTRODUCTION

Most animal species spend some part of their lives aggregated together in groups, and many benefits have been proposed and tested for this behaviour ^1,2^. For prey species, grouping behaviour can offer protection from predators through both the dilution of individual risk if an attack occurs ^3–5^ and an increase in the chance of successfully detecting an approaching predator due to the combined vigilance effort of the group ^5–7^, along with other anti-predator advantages of grouping behaviour such as synchronising activity to dilute risk ^8–11^. If an animal is being actively vigilant, it may be unable to conduct (or less efficient at) other important behaviours (like foraging or resting) at the same time (*e.g.* ^12^). Group membership means that vigilance can be pooled among the group members, which could mean that each individual can spend less time being vigilant and more time conducting other fitness-enhancing behaviours. A rich body of theory and research has explored how group size and individual vigilance effort are related ^13–17^, with much of it focussing on the prediction that individual vigilance effort will decrease as the group becomes larger. This prediction requires each individual to show a trade-off between vigilance and other behaviours, where being actively vigilant either cannot occur at the same time as other behaviours, or leads to a reduction in the efficiency of other behaviours that are conducted at the same time as being vigilant.

Vigilance is usually assumed to be occurring when an animal is actively scanning its surrounding environment with its head upwards, although there is no obvious consensus in how vigilance is defined in any particular species (see ^18^ for discussion of this problem in studies on primates). Although scanning behaviour is likely to stop an animal from actively collecting food, this head-up activity may not completely interfere with simultaneously conducted behaviours, such as chewing or social interaction. If a behaviour that is recorded as vigilance allows an individual to do other things at the same time, then we may be falsely assuming that this behaviour incurs the time and attention costs that are associated with vigilance ^19^. Without careful experimentation, it is difficult to assess how much of an individual’s attention is devoted to vigilance when we observe scanning or other forms of vigilance-like behaviour, which may add to the huge variation (*e.g*. ^13^) in whether a study demonstrates that individual vigilance is related to group size or not.

Although it is difficult to define exactly when an individual is being vigilant, we may instead be able to define when it is *not* able to be vigilant. Blinking (the temporary closure of both eyes, involving movements of the eyelids ^20^) is a good example of an activity where an individual is momentarily unable to visually scan the environment. It is an essential maintenance behaviour to keep the eyes moist and clean ^21^, and is conducted tens of times every minute in some species of diurnal mammals ^22–24^ and birds ^25^. Although a blink takes only a fraction of a second, the sum of this loss of visual information over multiple blinks could be substantial for the individual. In humans, spontaneous blinking is accompanied by attentional suppression, where the individual experiences a blackout in visual attention for the duration of the blink, meaning that there is no awareness of the temporary blindness and lack of visual information whilst the blinking is occurring ^26,27^. Blinking suppresses activity in both the visual cortex and other areas of the brain that are associated with awareness of environmental change ^28^. If we assume that other animals show similar attentional suppression, then they are essentially blind and unaware of changes in their visual environment during each blink, which in turn means that they cannot be visually vigilant for predators. Even if they remain vigilant for olfactory and auditory cues during a blink, the loss of visual information will reduce the efficiency and timing of an animal’s response to an approaching predator.

An individual’s blink rate therefore presents a trade-off between the physiological benefits of blinking and the loss of visual information during the blink ^21^. If an animal needs to dedicate more time to vigilance in a risky environment, then it has to reduce or suppress blinking to accommodate this increased vigilance. This is anecdotally demonstrated in American crows *Corvus brachyrhynchos*, which reduce their blink rates when looking at potentially dangerous stimuli ^29^, in horses *Equus caballus*, which decrease their spontaneous blink rate in response to stress-inducing stimuli ^30^, and in grackles *Quiscalus mexicanus*, which inhibit their blinking when observing human faces in varying orientations ^31^. This link between blink rate and vigilance implies that blink rate will also be related to group size. As group size increases, theory predicts that individual vigilance can be reduced ^5^, and so any requirement to suppress blinking will be relaxed. Blink rate may therefore increase with an increase in group size. Evidence supporting this is anecdotal: a comparison of chickens *Gallus gallus* feeding solitarily or in pairs showed an increase in blink rate and proportion of time spent blinking in the group-feeding birds ^32^, while a comparison of the blink rates of olive baboons *Papio anubis* ^33^ showed individuals in a small group blinked less than those in a large group (although the two groups were studied in different years). Here, we test this hypothesis by observing the blink rates in a captive herd of group-living red deer *Cervus elaphus*. Red deer are well-studied ungulates that exist both in wild populations and in managed captivity ^34–36^, and are a sister species to wapiti (or North American elk, *C. canadensis*), where considerable work has been conducted exploring how vigilance behaviour is mediated by natural predator presence (*e.g.* ^37–44^). Given that vigilance has been shown in wapiti to be related to the sex and age of an individual ^37,44^, we included these individual characteristics in our analysis.

## METHODS

### Study area, time and subjects

This observational study was conducted on the herd of red deer within the 40.5 hectare deer park in Ashton Court Estate, Bristol, England (51.4440° N, 2.6378° W), which is composed mainly of open grassland, with scattered forestry and a small area of water. The herd, managed by Bristol City Council, consists of *c*. 110 individuals of varying age and sex, who appear to mix freely. The enclosure is open to the public outside of the rutting season, so the deer are habituated to both dogs, humans, motor vehicles and occasional horses, and may be able to hear the vocalisations of a nearby (but separately fenced) captive herd of fallow deer *Dama dama* ^45^. Our observations were conducted over five days during the rutting season; observations were restricted between 1200-1630 h so they were outside of the dawn and dusk peaks of regular rutting activity ^35^.

### Behavioural sampling and observations

A random individual was selected as described in previous research on this herd ^46^. The selected individual was observed and dichotomously aged (mature or young) and sexed (male or female). The individual was sexed by the presence of antlers, as after one year of age only males have antlers ^47^. Age was identified by an individual’s size and morphology (larger individuals were older). If the individual was observed suckling, it was discarded and randomisation was repeated, as young individuals are hard to sex and exhibit behaviours uncommon to the rest of the herd ^35^. A count of the total number of young/mature males and females, along with suckling young, was made on three different days. The rounded averages of these five demographic classes were calculated, and used in the pseudoreplication analyses presented.

Prior to the study, the observers (ZWR and JHR, who both conducted the measurements described) were trained in identifying the recorded behaviours, and pilot trials ensured repeatability of measurements. A blink was defined as a rapid full closure of the eye. Group size was arbitrarily defined as the number of individuals aggregated no more than five body lengths away from at least one member of the group containing the focal deer, meaning a measured group was composed of individuals associated by a chain rule of association (see ^48^ for discussion of defining groups by associated neighbours within arbitrary distances). Before starting any set of observations, the observers waited 10 minutes at the site to habituate the deer. Observations were conducted approximately 10-100 m away from focal individuals using a 30× zoom spotting scope (Avian ED82 Magnesium Scope). An observation for a single selected individual was recorded for a maximum of 10 minutes. At the start of each minute the group size was counted by one observer, with blink rate (blinks per minute) being continuously recorded for each minute by the other observer. If the deer’s eye(s) were obstructed or there was a human/animal disturbance the observations were stopped with the current minute’s measures being disregarded. 75 observations of randomly selected individuals were observed using this method (and the analysis below describes how we controlled for potential pseudoreplication due to repeated samples potentially being taken using the same individual).

### Statistical analysis

Data were recorded as blinks per minute with group size recorded for that minute. Individuals were recorded a mean of 4.9 (± 2.0 SD) minutes before the data collection had to be discontinued due to the observer’s view of the eyes being obstructed. For each individual, we calculated the mean number of blinks in a minute, and the mean group size per minute. To compensate for any unevenness caused by some mean values being based on more observations than others, we conducted the same analyses using just the first minute of data for all individuals (see Supplementary Material 1 and 2): these data gave qualitatively similar results to the analysis involving mean group sizes, and are not discussed further.

Using *R* 4.0.3 ^49^, we constructed a linear mixed effects model where the natural logarithm of mean blink rate was described by the natural logarithm of group size, and the maturity and sex of the focal individual, including the observation date as a random effect. Logarithms were used to satisfy model assumptions of normalised residuals. A full model including interactions was initially considered, but no interaction terms were significant and so the basic additive model with the three explanatory variables was used. The Supplementary Information includes annotated *R* code, alongside the full dataset.

Although attempts were made to avoid replicating observations on the same individual during an observational day, it is likely that the 75 datapoints include some repeat observations of the same individual, given that the herd had 97 observable individuals (of which there were estimated to be 11 mature males, 23 young males, 26 young females and 37 mature females; 9 nursing young were also observed, but not included in the analysis), and individuals within each class could not be individually identified accurately. To explore potential effects of this pseudoreplication, we conducted simulations where identities were randomly allocated to individuals in each age class, and all but one datapoint for each ‘assumed individual’ was removed from the dataset (code presented in the Supplementary Information). A linear model was then run on this filtered subsample of the dataset, harvesting the significance value. By conducting 100,000 independent repeats of this resampling, we could identify how likely our dataset was to give a significant result (assuming significance was set at *p* = 0.05) if we had only sampled each individual once.

## RESULTS

Blink rate increased with group size (*t*_67_ = 11.38, *p* < 0.001; figure 1), and adults blinked more than young deer (*t*_67_ = 2.11, *p* = 0.038; figure 1). There was no relationship between sex and blinking (*t*_67_ = 0.35, *p* = 0.727). Considering potential pseudoreplication of unidentifiable individuals, resampling showed that group size remained significant (at *p* < 0.001) in 100% of simulations. There was a significant (at *p* < 0.05) effect of sex in only 0.17% of simulations, whilst maturity was significant in 40.73% of simulations. So, the effect of maturity observed could potentially have been an effect of resampling the same individuals, but the increasing blink rate in response to increasing group size is unlikely to have been affected by any pseudoreplication caused by repeated sampling of the same individual.

**Figure 1.**
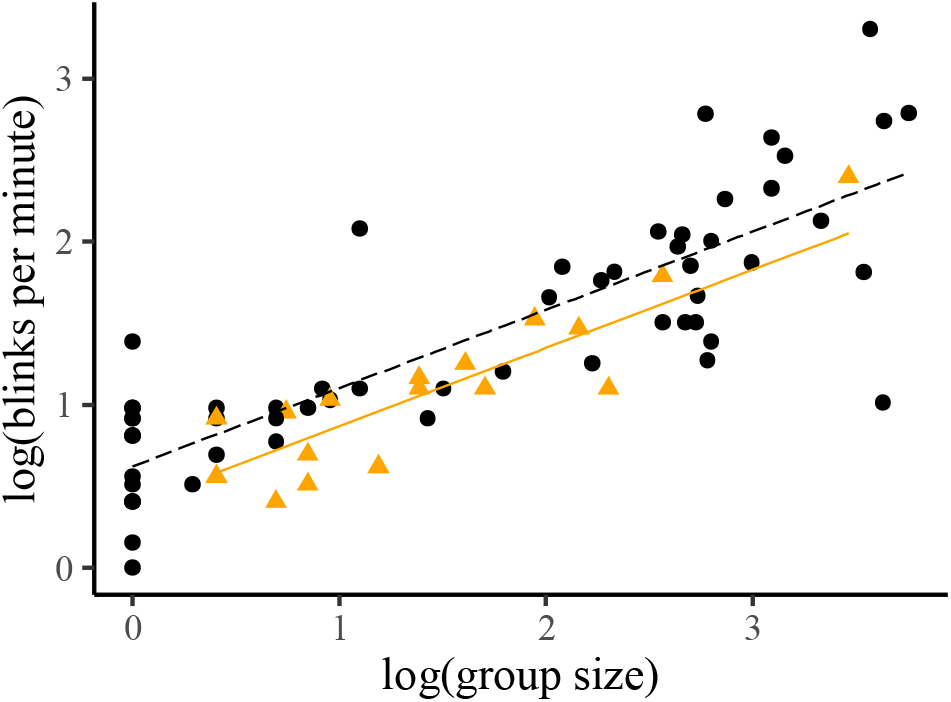
Scatterplot showing that blink rate increases with group size and maturity (where orange triangles and the fitted solid line represent young individuals, and black circles and the fitted dashed line represent adults).

## DISCUSSION

Our results demonstrate that blinking increases as group size increases. Given that blinking interferes with vigilance behaviour, and that individual vigilance is predicted to decrease as groups get larger, this supports our argument that the blink rate represents a trade-off between gaining visual information through vigilance and the physiological benefits of blinking. We note that these results are only correlational, and we suggest that a link between vigilance and blinking could be demonstrated with suitable experimental manipulation of perceived risk (*e.g.* ^8,50–52^).

We argued earlier that observed behaviours that are typically recorded as vigilance (such as holding a head upright or active scanning – see ^53,54^ for discussion) may not be conducted solely for vigilance (assuming that we are only considering visual vigilance, acknowledging that prey species will also be relying on auditory and olfactory information which will not be interrupted by eye closure). Similarly, blinking may not solely be a maintenance behaviour that is traded-off against being able to collect visual information. Blinking may include a social element, as rhesus macaques *Macaca mulatta* are able entrain their blink rate in response to social cues, coordinating their blinking with partners that they were interacting with ^55^. Our results suggest that the proportion of scanning time that an individual red deer allocates to vigilance is related to the size of its immediate group, but we should acknowledge that scanning behaviour may be influenced by social behaviour as well as vigilance. Studies of wild wapiti showed that young individuals were less vigilant than older ones ^37,42^, and suggest this may be because the younger individuals are unlikely to be able to outrun a predator if one appears. If the young in our study are following this behavioural pattern, this also suggests that blinking may not be completely correlated with vigilance level, as the younger deer should have had higher blink rates when compared to mature adults if the young were being less vigilant. Young deer lack experience of potential threats, may have different nutritional requirements and schedules to adults, and may not move their heads in a similar manner to adults, all of which could cause a difference between the age classes. This result may also reflect some form of social signalling between adults, although we note that we did not see a difference between males and females (which contradicts results from wapiti suggesting that vigilance patterns may be sex- and age-determined ^37,42^). Other aspects of social behaviour may also be important for determining blink rates, such as position within the group (echoing the vigilance changes seen in wapiti, where individuals on the outside of a group tend to be more vigilant ^42,43^). Theory predicts that individuals on the outside of the group should be more vigilant than those in the middle, and anecdotal evidence from olive baboons suggests that peripheral individuals may blink less ^33^. It may be possible to test many of the standard predictions connecting vigilance and group size (e.g. ^13–16^) using blink rate as a proxy for vigilance.

Observational studies on free-ranging wild wapiti living in national parks with varying levels of natural predation have variously found no effect of group size on vigilance (measured as heads-up scanning behaviour) in males and females ^39,40,42,43^, females alone ^44^, and one study ^37^ saw no significant relationship in males or breeding females and a decrease in vigilance with respect to increasing group size in non-breeding females. Model comparison ^41^ suggests that wapiti vigilance may show a non-linear relationship with herd size, where vigilance increases with small numbers, and then decreases after the herd has reached an intermediate size, whilst position in the herd was less important. These results suggest that wapiti do not normally show a decrease in individual vigilance in response to increasing group size, as would be predicted by standard theory (*e.g.* ^13–16^), which in turn either suggests that this predicted relationship does not occur, or that standard measures of vigilance using assays of head-up scanning behaviour may not be suitable for quantifying vigilance, as we have argued above. We did not assay heads-up scanning behaviour alongside the blinking behaviour measured here, and a sensible next step would be to measure both simultaneously, to assess whether red deer conform with wild wapiti in showing a similar lack of correlation between head-up scanning vigilance and group size.

It could be argued that our semi-captive deer population is not suitable for assaying something which is considered an anti-predator behaviour, as the population lives in a relatively benign managed environment, with no natural predators present. In wild wapiti, the very real risk of predation by wolves (which have been recently reintroduced) or other carnivores has a measurable impact on behaviour, affecting both observable behaviour and use of space in the environment ^39^. Wapiti may respond to patchiness of predation risk in the environment by choosing where they spend their time, which in turn can mediate whether their diet changes ^56^, and whether they show a stress response (or not, *e.g.* ^38^).

However, there is evidence to suggest that red deer living in what we consider to be a predator-free environment do nonetheless show responses to anthropogenic cues and features in the environment that they may be treating in the same way as a predator cue. Free-ranging wapiti were shown to increase vigilance and their likelihood of flight behaviour in response to vehicle presence ^57,58^, and vigilance behaviour has been observed as being more influenced by both traffic and transport-related infrastructure than by predator presence ^59^. Other evidence suggests that red deer mediate their behaviour greatly in response to human presence. Vigilance was more likely in areas with human disturbance in the Scottish Highlands ^60^. Wild free-ranging individuals in southern Germany were shown to avoid areas with high human recreational presence during the day ^61^, and a comparison of free-ranging populations from different regions of Poland with differing natural predation levels showed that faecal glucocorticoid concentrations were lowest and least variable in high predation areas, and that concentrations indicating high stress were instead likely to be linked to the level of anthropogenic disturbance that they were experiencing ^62^. Similarly, measurement of faecal glucocorticoids in two herds of semi-wild red deer living in parkland in England similar to the current study demonstrated that assayed stress was higher on days with higher visitor numbers ^63^. All this evidence suggests that red deer confined to a small area with constant disturbance by both pedestrian visitors and motor vehicles may well show stress responses and anti-predator behaviour that may be differently expressed when compared to deer living in an undisturbed wild environment with natural predators present. It would be interesting to examine whether these different stressors cause different responses, to disentangle whether the blinking response that we see is coupled with ‘head-up’ scanning vigilance, or whether scanning is not related to group size as is seen in the multiple wild wapiti studies described above. This could be done by considering red deer in environments where wolves are present, or else by manipulating ‘natural’ predator cues such as by adding wolf urine to the environment (^64^, although red deer quickly acclimatise to this manipulation ^65^). However, it is also useful to question whether the lack of ‘natural’ predators is going to stop the deer performing the vigilance behaviours that they have evolved. As a relevant example, we could consider Père David’s deer *Elaphurus davidianus*; this endangered species has only existed in a managed, predator-free environment for over a thousand years, but nonetheless shows distinct group size-related vigilance behaviour in response to human disturbance ^66^, demonstrating that vigilance behaviour does not need wolves or tigers to be visible in a species.

Blink rate may also be influenced by factors other than the size of the group, including rainfall and wind (which have been shown to influence blink rate in captive grackles ^67,68^). Similarly, the behaviour that an individual conducts simultaneously to the blink may be important. Experiments in humans suggest a mechanism controlling the timing of blinks, which occurs to minimise the chance of missing crucial information ^21,69–72^, with evidence of similar behaviours in rhesus macaques *Macaca mulatta* ^55^. Peafowl *Pavo cristatus* also time their blinks to coincide with gaze shifts ^73^ while grackles blink less during flight behaviours ^74^ and chickens blink more when feeding when compared to scanning ^32^, all minimising the time where visual information cannot be collected from the environment. Therefore, if an individual is moving, the timing and frequency of its blinks may reflect this movement, and it may therefore be sensible to assay blinking in response to group size in resting deer groups, which would not be undergoing head and body movements that could confound the measure of blinking that is recorded. Similarly, animals in different attentional states may change their frequency of blinking, such as during sleep in herring gulls *Larus argentatus* ^75^.

Our results suggest that the measurement of blinking presents a simple and non-invasive technique for observing attention that can be conducted remotely. Although we conducted our sampling in the field, this could be done using video footage. Being able to analyse video footage means that information about blink duration can also be collected, and previous studies have demonstrated that this additional metric can also vary between individuals and species ^23,24,32,33,67,68^, and may increase in relation to group size ^32^. From a field observation perspective, being able to zoom in on detail may also be extremely useful if the observed individuals are a long distance away (as we acknowledge we were lucky with being able to get within 100m of our sample animals). Given that blinking has been shown to decrease under stressful conditions ^29,30^, this simple technique could help us to understand the welfare requirements of managed animals that normally live in social groups.

## Supporting information

Raw data

## ACKNOWLEDGEMENTS

ZWR and JHR conducted this work as their final year BSc projects. We thank the School of Biological Sciences teaching lab team for help with equipment, and several reviewers whose comments greatly improved the manuscript. Ethical permission to conduct the study was given by the University of Bristol Animal Welfare and Experimental Review Board (#UB/16/062). SAR was supported a Natural Environment Research Council (UK) standard grant (NE/P012639/1).

## AUTHOR CONTRIBUTIONS

ZWR: Conceptualisation, experimental design, data collection, drafting, and editing. JHR: Conceptualisation, experimental design, data collection, drafting, and editing. SAR: Conceptualisation, supervision, experimental design, analysis, original draft, and editing.

## SUPPLEMENTARY INFORMATION

**Supplementary Information 1**. Text file ‘blinkdata.txt’, presenting the data used as a space-delimited plain text file. ‘blinks’ denotes number of mean number of blinks per minute for a focal individual; ‘groupsize’ records the mean group size for the focal individual; ‘deerdata’ denotes whether the focal individual is a mature female (MF), mature male (MM), young female (YF), or young male (YM); ‘sex’ denotes whether the focal individual is male (m) or female (f); ‘age’ denotes whether the focal individual is mature (m) or young (y); and ‘day’ denotes the day the observation was made.

**Supplementary Information 2**. Text file ‘deer blinking analysis code.R’, presenting the annotated *R* code used for all analyses described in the manuscript.

