## Supplementary material for "Red deer *Cervus elaphus* blink more in larger groups": R code

library(ggplot2)
library(nlme)

# Reading the dataset
blink <- read.table("blinkdata.txt", header=T)


lm0 <- lme(log(blinks)~log(groupsize)*sex*age, data=blink, random=~1|day, method="ML")
summary(lm0)
anova(lm0)
#some non-sig terms - the three-way interaction is non-sig, so trying model without...
lm1 <- lme(log(blinks)~log(groupsize)*sex*age - log(groupsize):sex:age, data=blink, random=~1|day, method="ML")
summary(lm1)
anova(lm1)
#and comparing
anova(lm0, lm1)
#although slightly higher AIC for full model, no significant diff between two, so sticking with most parsimonious (lm1), which removes the three-way interaction

#none of the two-way interactions in lm1 are significant, so running a batch of models to check whether single or multiple removal has an impact:
lm2 <- lme(log(blinks)~log(groupsize)*sex*age - log(groupsize):sex:age - log(groupsize):sex , data=blink, random=~1|day, method="ML")
lm3 <- lme(log(blinks)~log(groupsize)*sex*age - log(groupsize):sex:age - log(groupsize):age , data=blink, random=~1|day, method="ML")
lm4 <- lme(log(blinks)~log(groupsize)*sex*age - log(groupsize):sex:age - sex:age , data=blink, random=~1|day, method="ML")
lm5 <- lme(log(blinks)~log(groupsize) + sex + age + log(groupsize):sex, data=blink, random=~1|day, method="ML")
lm6 <- lme(log(blinks)~log(groupsize) + sex + age + log(groupsize):age, data=blink, random=~1|day, method="ML")
lm7 <- lme(log(blinks)~log(groupsize) + sex + age + sex:age, data=blink, random=~1|day, method="ML")
lm8 <- lme(log(blinks)~log(groupsize) + sex + age, data=blink, random=~1|day, method="ML")
anova(lm2,lm3,lm4, lm5,lm6, lm7, lm8)
#get
#Model df      AIC      BIC    logLik   Test   L.Ratio p-value
#lm2     1  8 79.46135 98.00125 -31.73067                         
#lm3     2  8 77.24314 95.78305 -30.62157                         
#lm4     3  8 78.07531 96.61522 -31.03766                         
#lm5     4  7 77.11972 93.34214 -31.55986 3 vs 4 1.0444109  0.3068
#lm6     5  7 77.46399 93.68641 -31.73200                         
#lm7     6  7 77.78374 94.00616 -31.89187                         
#lm8     7  6 76.05612 89.96105 -32.02806 6 vs 7 0.2723803  0.6017
#where the model without interactions (lm8) is the most parsimonious, and has the lowest AIC. So can safely ignore the interaction terms


#model described in paper, with all variables but no interactions
lm8 <- lme(log(blinks)~log(groupsize) + sex + age, data=blink, random=~1|day, method="ML")
summary(lm8)
anova(lm8) 
plot(lm8) #all model assumptions look like they're satisfied here

# This model gives the following parameter estimates
#
#Linear mixed-effects model fit by maximum likelihood
#Data: blink 
#AIC      BIC    logLik
#76.05612 89.96105 -32.02806
#
#Random effects:
#  Formula: ~1 | day
#(Intercept)  Residual
#StdDev: 7.923612e-06 0.3708722
#
#Fixed effects: log(blinks) ~ log(groupsize) + sex + age 
#Value  Std.Error DF   t-value p-value
#(Intercept)     0.6119165 0.09512963 67  6.432450  0.0000
#log(groupsize)  0.4796761 0.04215537 67 11.378767  0.0000
#sexm            0.0439175 0.12522903 67  0.350697  0.7269
#agey           -0.2338815 0.11061626 67 -2.114350  0.0382
#Correlation: 
#  (Intr) lg(gr) sexm  
#log(groupsize) -0.828              
#sexm           -0.590  0.509       
#agey           -0.089 -0.111 -0.307
#
#Standardized Within-Group Residuals:
#  Min          Q1         Med          Q3         Max 
#-3.61853086 -0.63884448  0.09821871  0.58807522  2.64489355 
#
#Number of Observations: 75
#Number of Groups: 5 
#
#numDF denDF  F-value p-value
#(Intercept)        1    67 903.3826  <.0001
#log(groupsize)     1    67 172.9383  <.0001
#sex                1    67   0.0978  0.7554
#age                1    67   4.4705  0.0382


#Code for producing the plot.
#This ggplot2 code produces a plot of log(group size) as the explanatory axis, and log(blin rate) as the dependent
#The adult data are plotted as black circles, and the juvenile data as orange triangles.
#The predicted model lines are fitted for the two age classes in the given colours.

log_smallest_group_for_adult <- log(min(subset(blink,age=="m")$groupsize))
log_largest_group_for_adult <- log(max(subset(blink,age=="m")$groupsize))
log_smallest_group_for_juvenile <- log(min(subset(blink,age=="y")$groupsize))
log_largest_group_for_juvenile <- log(max(subset(blink,age=="y")$groupsize))
eq1 = function(x)coef(lm1)[1]+coef(lm1)[2]*x+(17/75)*coef(lm1)[3]
eq2 = function(x)coef(lm1)[1]+coef(lm1)[2]*x+(17/75)*coef(lm1)[3]+coef(lm1)[4]
p1 <- ggplot(blink, aes(log(groupsize),log(blinks)))
p1 + geom_point(aes(colour = age,shape=age),size=3,show.legend=FALSE)+
  scale_color_manual(values=c("black", "orange"))+
  stat_function(fun=eq1,geom="line",xlim=c(log_smallest_group_for_adult,log_largest_group_for_adult),linetype=5)+
  stat_function(fun=eq2,geom="line",colour="orange",xlim=c(log_smallest_group_for_juvenile,log_largest_group_for_juvenile))+
  theme_classic(base_size=20, base_family="Times")+
  labs(x="log(group size)",y="log(blinks per minute)")
# (These settings work nicely if you export the figure as a pdf sized 4.5" x 6")


#To CHECK, SIMILAR TEST DONE WITH JUST FIRST MINUTE DATA (BLINKS IN FIRST MINUTE VERSUS GROUP SIZE IN FIRST MINUTE)
blink1min <- read.table("firstminutedata_WITH_DATES.txt", header=T)
lm1min <- lme(log(blinks)~log(groupsize) + sex + age, data=blink1min, random=~1|day, method="ML")
anova(lm1min)
plot(log(blink1min$blinks) ~ log(blink1min$groupsize), xlab=c("ln(group size)"), ylab=c("ln(blinks per minute)"))
plot(lm1min)
#GIVES QUALITATIVELY SAME RESULTS AND TRENDS


#TESTING FOR PSEUDOREPLICATION, WITH CLASSES AS DEFINED (106 indivs of which 9 too young, 11 mm, 23 ym, 26 yf and 37 mf)
blink$identity <- NA
collected_p_groupsize <- NULL
collected_p_sex <- NULL
collected_p_age <- NULL
for (j in 1:10000) {
  assigned_identity <- NA
  for (i in 1:48) assigned_identity[i] <- sample(1:37,1)
  assigned_identity <- sort(assigned_identity)
  for (i in 47:1) if (assigned_identity[i] == assigned_identity[i+1]) { assigned_identity[i+1] <- 0 }
  for (i in 1:48) if (assigned_identity[i] > 0) assigned_identity[i] <- 1
  ordering <- sample(1:48)
  for (i in 1:48) blink$identity[ordering[i]] <- assigned_identity[i]

  assigned_identity <- NA
  for (i in 1:9) assigned_identity[i] <- sample(1:26,1)
  assigned_identity <- sort(assigned_identity)
  for (i in 8:1) if (assigned_identity[i] == assigned_identity[i+1]) { assigned_identity[i+1] <- 0 }
  for (i in 1:9) if (assigned_identity[i] > 0) assigned_identity[i] <- 1
  ordering <- sample(49:57)
  for (i in 1:9) blink$identity[ordering[i]] <- assigned_identity[i]
  
  assigned_identity <- NA
  for (i in 1:10) assigned_identity[i] <- sample(1:11,1)
  assigned_identity <- sort(assigned_identity)
  for (i in 9:1) if (assigned_identity[i] == assigned_identity[i+1]) { assigned_identity[i+1] <- 0 }
  for (i in 1:10) if (assigned_identity[i] > 0) assigned_identity[i] <- 1
  ordering <- sample(58:67)
  for (i in 1:10) blink$identity[ordering[i]] <- assigned_identity[i]
  
  assigned_identity <- NA
  for (i in 1:8) assigned_identity[i] <- sample(1:23,1)
  assigned_identity <- sort(assigned_identity)
  for (i in 7:1) if (assigned_identity[i] == assigned_identity[i+1]) { assigned_identity[i+1] <- 0 }
  for (i in 1:8) if (assigned_identity[i] > 0) assigned_identity[i] <- 1
  ordering <- sample(68:75)
  for (i in 1:8) blink$identity[ordering[i]] <- assigned_identity[i]
  
  subsampled_set <- subset(blink, identity == 1)
#  lm_temp <- lm(log(subsampled_set$blinks) ~ log(subsampled_set$groupsize) + subsampled_set$sex + subsampled_set$age)
  lm_temp <- lme(log(blinks)~log(groupsize) + sex + age, data=subsampled_set, random=~1|day, method="ML")  

  collected_p_groupsize <- c(collected_p_groupsize, anova(lm_temp)[,4][2])
  collected_p_sex <- c(collected_p_sex, anova(lm_temp)[,4][3])
  collected_p_age <- c(collected_p_age, anova(lm_temp)[,4][4])
}  
plot(sort(collected_p_groupsize))
plot(sort(collected_p_sex))
plot(sort(collected_p_age))
(10000-length(which(collected_p_groupsize>0.05)))/10000
(10000-length(which(collected_p_sex>0.05)))/10000
(10000-length(which(collected_p_age>0.05)))/10000


plot(blink$groupsize~blink$sex)
t.test(blink$groupsize~blink$sex)


femaleblink <- read.table("blinkdata_female_only.txt", header=T)

lmf <- lm(log(blink$blinks) ~ log(blink$groupsize)  + blink$age)

anova(lmf)
plot(log(blink$blinks) ~ log(blink$groupsize))
plot(lmf)


#SECOND ATTEMPT, WITH CLASSES AS DEFINED (106 indivs of which 9 too young, 11 mm, 23 ym, 26 yf and 37 mf)
femaleblink$identity <- NA
collected_p_groupsize <- NULL
collected_p_sex <- NULL
collected_p_age <- NULL
for (j in 1:10000) {
  assigned_identity <- NA
  for (i in 1:48) assigned_identity[i] <- sample(1:37,1)
  assigned_identity <- sort(assigned_identity)
  for (i in 47:1) if (assigned_identity[i] == assigned_identity[i+1]) { assigned_identity[i+1] <- 0 }
  for (i in 1:48) if (assigned_identity[i] > 0) assigned_identity[i] <- 1
  ordering <- sample(1:48)
  for (i in 1:48) femaleblink$identity[ordering[i]] <- assigned_identity[i]
  
  assigned_identity <- NA
  for (i in 1:9) assigned_identity[i] <- sample(1:26,1)
  assigned_identity <- sort(assigned_identity)
  for (i in 8:1) if (assigned_identity[i] == assigned_identity[i+1]) { assigned_identity[i+1] <- 0 }
  for (i in 1:9) if (assigned_identity[i] > 0) assigned_identity[i] <- 1
  ordering <- sample(49:57)
  for (i in 1:9) femaleblink$identity[ordering[i]] <- assigned_identity[i]
  
  subsampled_set <- subset(femaleblink, identity == 1)
  lm_temp <- lm(log(subsampled_set$blinks) ~ log(subsampled_set$groupsize) + subsampled_set$age)
  
  collected_p_groupsize <- c(collected_p_groupsize, anova(lm_temp)[5][,1][1])
  collected_p_age <- c(collected_p_age, anova(lm_temp)[5][,1][2])
}  
plot(sort(collected_p_groupsize))
plot(sort(collected_p_age))
(10000-length(which(collected_p_groupsize>0.05)))/10000
(10000-length(which(collected_p_age>0.05)))/10000
plot(log(femaleblink$blinks)~log(femaleblink$groupsize))

plot(blink$groupsize~blink$sex)
t.test(blink$groupsize~blink$sex)
plot(blink$blinks~blink$age)
